# Viral genome-capsid core senses host environments to prime RNA release

**DOI:** 10.1101/2021.11.23.469659

**Authors:** Ranita Ramesh, Sean M. Braet, Varun Venkatakrishnan, Palur Venkata Raghuvamsi, Jonathan Chua Wei Bao, Jan K. Marzinek, Chao Wu, Ruimin Gao, Sek-Man Wong, Peter J. Bond, Ganesh S. Anand

## Abstract

Viruses are metastable macromolecular assemblies containing a nucleic acid core packaged by capsid proteins that are primed to disassemble in host-specific environments leading to genome release and replication. The mechanism of how viruses sense environmental changes associated with host entry to prime them for disassembly is unknown. We have applied a combination of mass spectrometry, cryo-EM, and simulation-assisted structure refinement to Turnip crinkle virus (TCV), which serves as a model non-enveloped icosahedral virus (Triangulation number = 3, 180 copies/icosahedron). Our results reveal genomic RNA tightly binds a subset of viral coat proteins to form a stable RNA-capsid core which undergoes conformational switching in response to host-specific environmental changes. These changes include: i) Depletion of Ca^2+^ which triggers viral particle expansion ii) Increase in osmolytes further disrupt interactions of outer coat proteins from the RNA-capsid core to promote complete viral disassembly. A cryo-EM structure of the expanded particle shows that RNA is asymmetrically extruded from a single 5-fold axis during disassembly. The genomic RNA:capsid protein interactions confer metastability to the TCV capsid and drive release of RNA from the disassembling virion within the plant host cell.

**AUTHOR SUMMARY:** RNA viruses including coronaviruses, dengue, influenza, and HIV are a significant threat to human health. These viral particles are finely tuned to undergo complex conformational changes that allow for response to varied environments. Turnip crinkle virus (TCV) serves as an excellent model for studying RNA virus dynamics. Since TCV is non-enveloped and has no post-translational modifications, we can specifically investigate the contributions of RNA to viral dynamics. Genomic RNA is not a passive entity but plays a crucial and previously uncharacterized role in viral disassembly. Our results reveal that the genomic RNA-capsid core serves as an environmental sensor and undergoes conformational switching in response to host cell conditions.

## INTRODUCTION

Icosahedral viruses are ubiquitous macromolecular assemblies, consisting of protein capsids encapsidating nucleic acid genomes. Capsids function to shield the viral genome and undergo programmed disassembly in favorable host environments in order to release the genome for replication inside the target host cell. Coat proteins that form the capsid thus function as metastable sensors of environmental conditions through reversible ‘breathing’ motions [1,2] and can be likened to delivery vehicles of genetic material transmitted from one host to another. Cryo-electron microscopy (cryo-EM) has provided high resolution snapshots of virus icosahedral geometries [3]. However, static structural snapshots cannot capture intrinsic breathing dynamics [4,5] of the virus particle, and largely report on more structured elements of the virus particle and offer only limited insights into the interior of the viral particle.

Whole viral particle dynamics are fundamental to understanding a virus’s structural responses to environmental changes [6]. Viruses are programmed to sense and respond to unique host specific physical perturbations such as changes in osmolyte concentration [7], pH [8] and temperature [9], or chemical perturbations such as changes in divalent cation concentrations [10]. Currently, viral recognition and responses to host environments is poorly understood. Here, we describe the intrinsic dynamics of virus particles and changes in conformation associated with host-specific environments *in vitro*.

We selected Turnip crinkle virus (TCV), a plant RNA icosahedral virus model for mapping changes in viral particle conformation and dynamics across varying plant host-specific environments. TCV offers numerous advantages: It is a simple non-enveloped virus lacking lipids and post-translational glycosylation modifications and consists entirely of 180 copies of a viral capsid coat protein (38 kDa) arranged in T=3 icosahedral symmetry and encapsulating a 4.1 kb ssRNA genome [11]. Each coat protein subunit consists of 3 domains – an N-terminal RNA binding (R) domain, a Shell (S) domain, and a C-terminal Protruding (P) domain. This basic unit consists of a trimer of 3 quasi equivalent conformations - A, B and C - that differ in their hinge angles between the S and P domains and folding of the connecting arm linking R and S domains [12]. Upon entry into a host plant cell, TCV undergoes preferential disassembly to release genomic RNA into the associated high osmolyte environment with concomitant depletion of divalent Ca^2+^ ions in the host cytosol (Fig 1). Release of viral RNA without complete disassembly has also been observed whereby chelation of divalent Ca^2+^ ions results in a large expansion in particle size (disassembly intermediate). Subsequent proteolytic processing of the expanded viral particle by host proteases facilitates ribosomal-mediated extrusion (striposomal release) of genomic RNA into the plant host [13].

**Figure 1.**
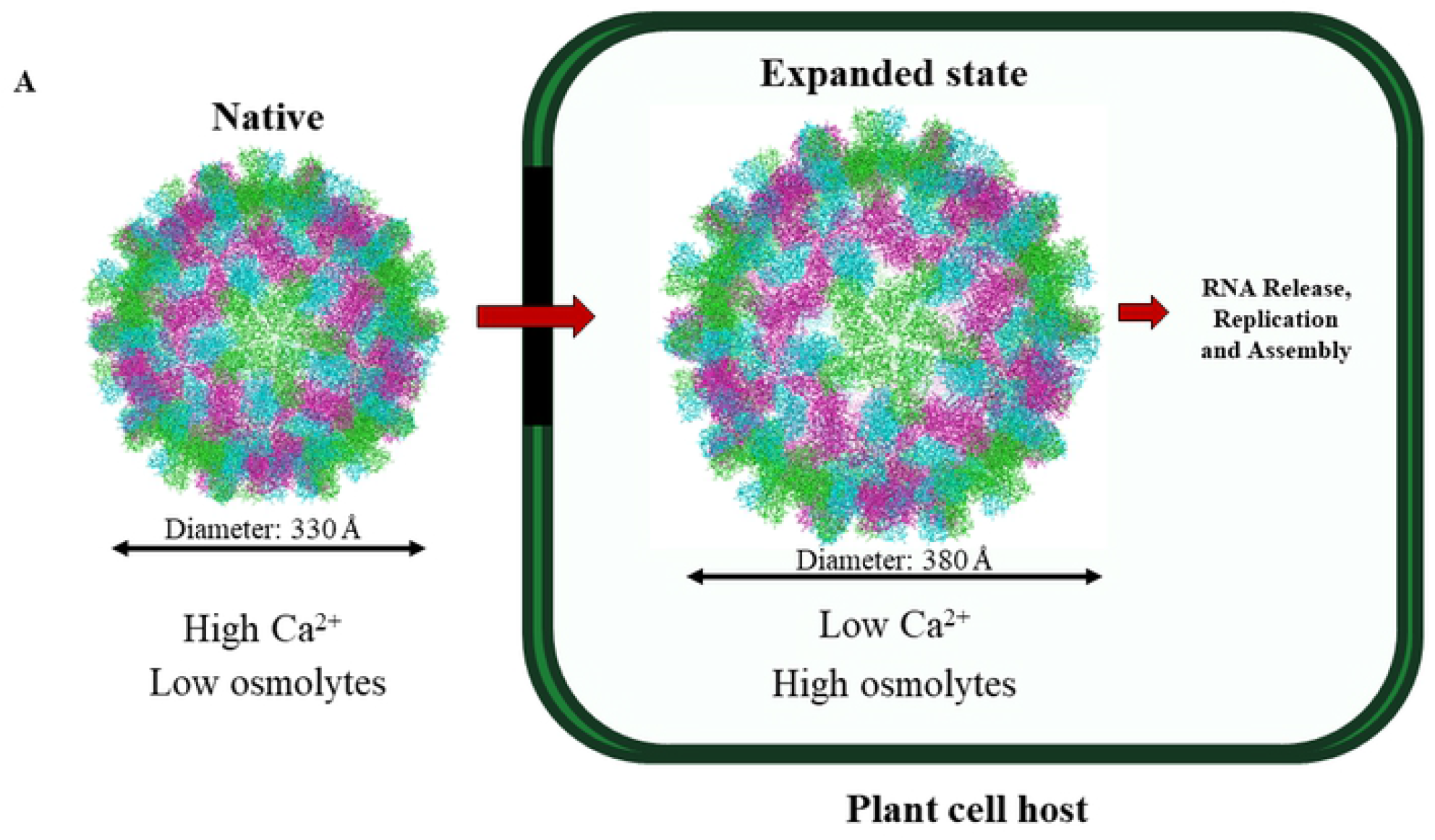
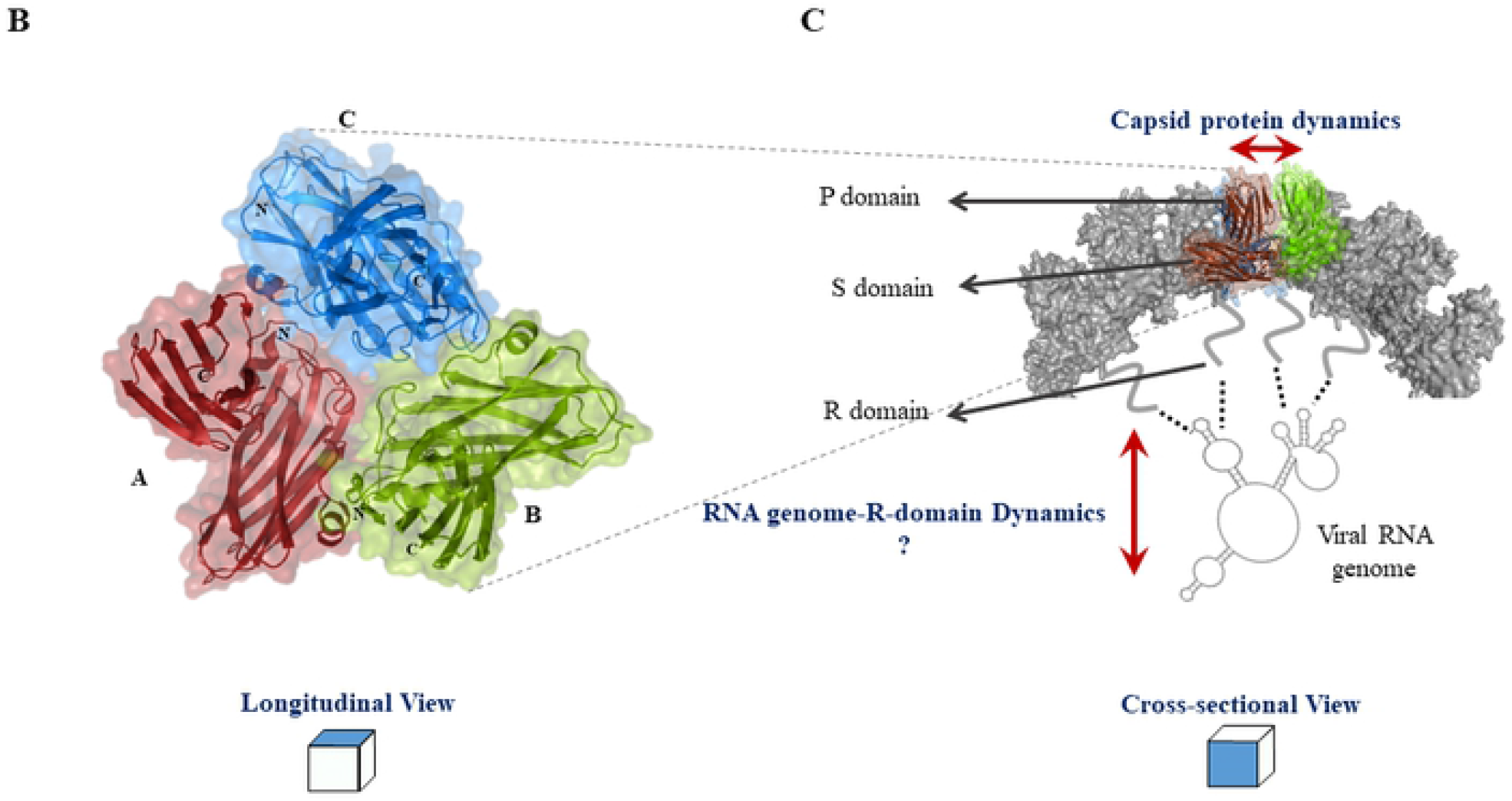
Changes in Turnip crinkle virus (TCV) particle upon host entry. **(A)** Environmental changes associated with passive entry of TCV into a plant host include decreases in intracellular Ca^2+^ and increased osmolality, which trigger disassembly of the TCV viral particle to release viral genomic RNA for replication. Complete viral disassembly is preceded by formation of a disassembly intermediate, the ‘Expanded’ state with a viral particle diameter of ~380 Å. Native (PDB ID: 3ZX8) and ‘Expanded’ (PDB ID: 3ZX9) TCV are in surface representation showing the 3 quasi-equivalent subunits of the TCV basic asymmetric trimeric unit (A – red, B – green, C – blue). (PDB ID of ‘Native TCV’ – 3ZX8, ‘Expanded TCV’ – 3ZX9) [13]. **(B)** TCV asymmetric unit with 3 quasi-equivalent conformations A, B and C in red, green and blue, respectively. **(C)** Cross-section of TCV depicting key interactions between the coat protein and viral genomic RNA. Three domains of the coat protein Protruding (P Domain), Shell (S Domain) and RNA Binding (R Domain). The contribution of R domain and genomic RNA interactions to TCV metastability are unknown.

Cryo-EM structures of the native virus and expanded disassembly intermediates have previously been solved at low resolution (11 and 18 Å respectively). These structures show a disordered arm in the A and B conformations and more folded structure in the C conformation. The A subunits form the 5-fold axis, while the B and C subunits form the 3-fold (or quasi 6-fold) axis. The arms of the C subunit form a beta annulus structure at the 3-fold axis that has been postulated to be involved in RNA binding [13].

To obtain additional insights into the interior of the virus particle, we have carried out orthogonal cryo-EM, simulation-assisted structure refinement, and amide hydrogen/deuterium exchange mass spectrometry (HDXMS) analysis of the different states of TCV. HDXMS is a powerful technique that can be used to identify regions involved in conformational changes within a protein, and within protein-protein, protein-lipid, protein-ligand or protein-nucleic acid interfaces by reporting on hydrogen bonding and solvent accessibility [14,15]. HDXMS also allows detection of multiple conformational populations in solution and has been combined with variable urea denaturation to estimate the relative strengths of the icosahedral viral assembly [16,17].

We report a high resolution (3.2 Å) map that clearly shows the asymmetry in the interior of the TCV particle and captures TCV in the process of releasing its genomic RNA into the host plant cell. While the icosahedral geometry can be clearly described at this resolution, the specific RNA-R domain binding interactions inside the whole virus particle are still unresolved and reflect a highly dynamic interior, corroborated by HDXMS. Our findings further reveal a highly disordered R-domain that is composed of genomic RNA-tightly bound (~5.7%) capsid protein. Significantly, a majority of the capsid protein (~94.3%) is only loosely bound to the genomic RNA-capsid core. The basis for the TCV virion assembly lies in a subset of the capsid protein population binding strongly to RNA to generate a ribonucleoprotein core capable of nucleating assembly of many more capsid protein units to generate an icosahedral particle. The intrinsic metastability of native TCV is explained by the interplay between the small subset of strong capsid RNA interactions holding together the RNA-capsid core and weaker capsid interactions maintaining the icosahedral geometry. Far from being a passive entity, the genomic RNA-capsid core complex is the ‘controller switch’ for virus assembly-disassembly transitions.

## RESULTS

### R domain of coat protein shows greatest relative deuterium exchange

HDXMS of TCV at t = 1, 10 and 30 min, covering the fast deuterium exchange kinetic timescales [18] was carried out as described in Materials and Methods. 61 peptides with high signal to noise spanning the entire TCV coat protein with 89.2% sequence coverage were obtained (S1 Fig and S1 Table).

Of the three domains, the R domain showed the greatest magnitude exchange while the S and P domains showed lower relative exchange overall, consistent with the R domain being the least structurally ordered domain (Fig 2A). Mass spectral envelopes of N-terminal peptides spanning the R domain upon deuterium exchange showed two distinct populations corresponding to a low exchanging (blue) and high exchanging (green) population respectively (Fig 2B). These were deconvolved using HDExaminer 3.0 as previously reported [19]. The low exchanging population showed no increases in exchange with time while the mass spectral envelope of the higher exchanging population shifted to the right in the deuteration times examined. Bimodal deuterium exchange profiles for all times (t = 1, 10 and 30 min) were observed for the 19 N-terminal fragment peptides and not observed for any of the peptides from the S or P domains (Fig 2C and 2D). We report peptide 66-89 as a reference peptide to quantitate the abundance of low and high exchanging populations in native TCV, which was consistent with other R domain peptides. A small (5.7%) fraction of the R domain in the native TCV virus particle with lower exchange (~6.2 Da) while a majority of the R domain (~94.3%) showed a higher exchange (~11.7 Da) (S2 Fig).

**Figure 2.**
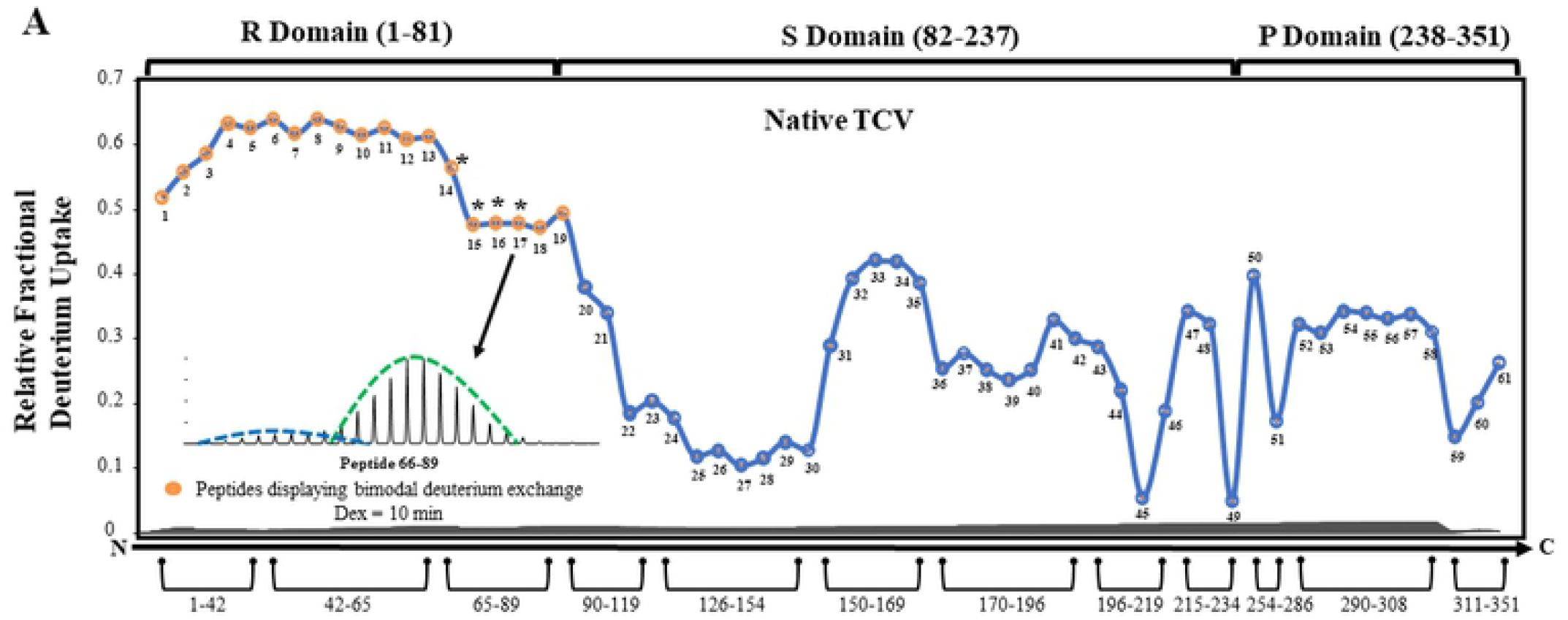
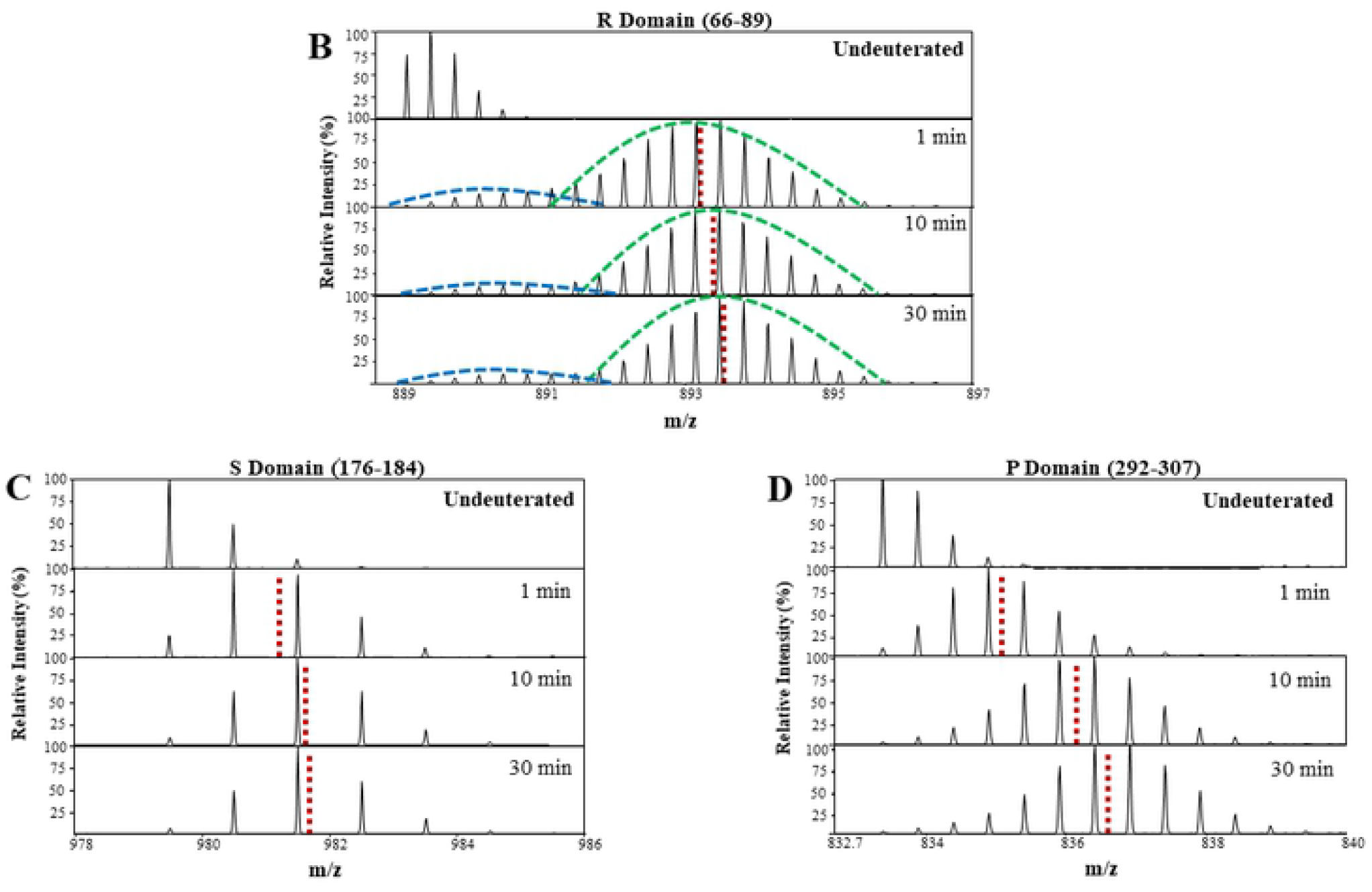
TCV R domain exists in an ensemble of two conformations. **(A)** Relative Deuterium fractional uptake (RFU) plot of pepsin fragment peptides of the TCV capsid protein (Native state) at Dex = 10 min for all peptides listed from the N to C-terminus (X-axis). Orange nodes denote peptides showing bimodal mass spectra for deuterium exchange while blue nodes represent peptides showing unimodal (binomial) spectra. ‘*’ indicates peptides spanning Tyr 66 (site of limited chymotrypsin proteolysis). Inset shows the mass spectral plot of peptide 66-89 (Dex = 10 min) with low exchanging population distribution outline in blue and high exchanging population distribution outlined in green. Mass spectral plots comparing the undeuterated and deuterated states after Dex = 1 min, 10 min, and 30 min for representative peptides from the **(B)** R domain - 66-89, **(C)** S domain - 176-184 and **(D)** P domain - 292-307. Red dotted lines indicate the centroid masses of each of the spectral envelopes.

### TCV expansion shows enhanced deuterium exchange at quasi 3-fold axes along with changes in R-domain ensemble behavior

To capture conformational changes associated with particle expansion, we next carried out HDXMS of expanded TCV which was generated by chelation of Ca^2+^ with EDTA. Differences in exchange between the expanded and native TCV particles are represented as a deuterium exchange difference plot (Fig 3A). Increases in exchange were observed predominantly in the S domain. These regions exhibiting increases in exchange greater than 0.5 D are indicated in red (Fig 3B). Increases in exchange observed at peptides spanning 170-176 and 196-208 of the S domain correspond to the region flanking the quasi 3-fold axis which shows increases in solvent accessibility, confirming particle expansion. This can be observed through shifts in the mass centroids of spectral envelopes from the native to the expanded state in representative peptide 150-159 (Fig 3D). Increases in exchange were additionally observed at the 3-fold and 5-fold axes, corroborating our previous results showing these loci as hotspots for TCV disassembly [16].

**Figure 3.**
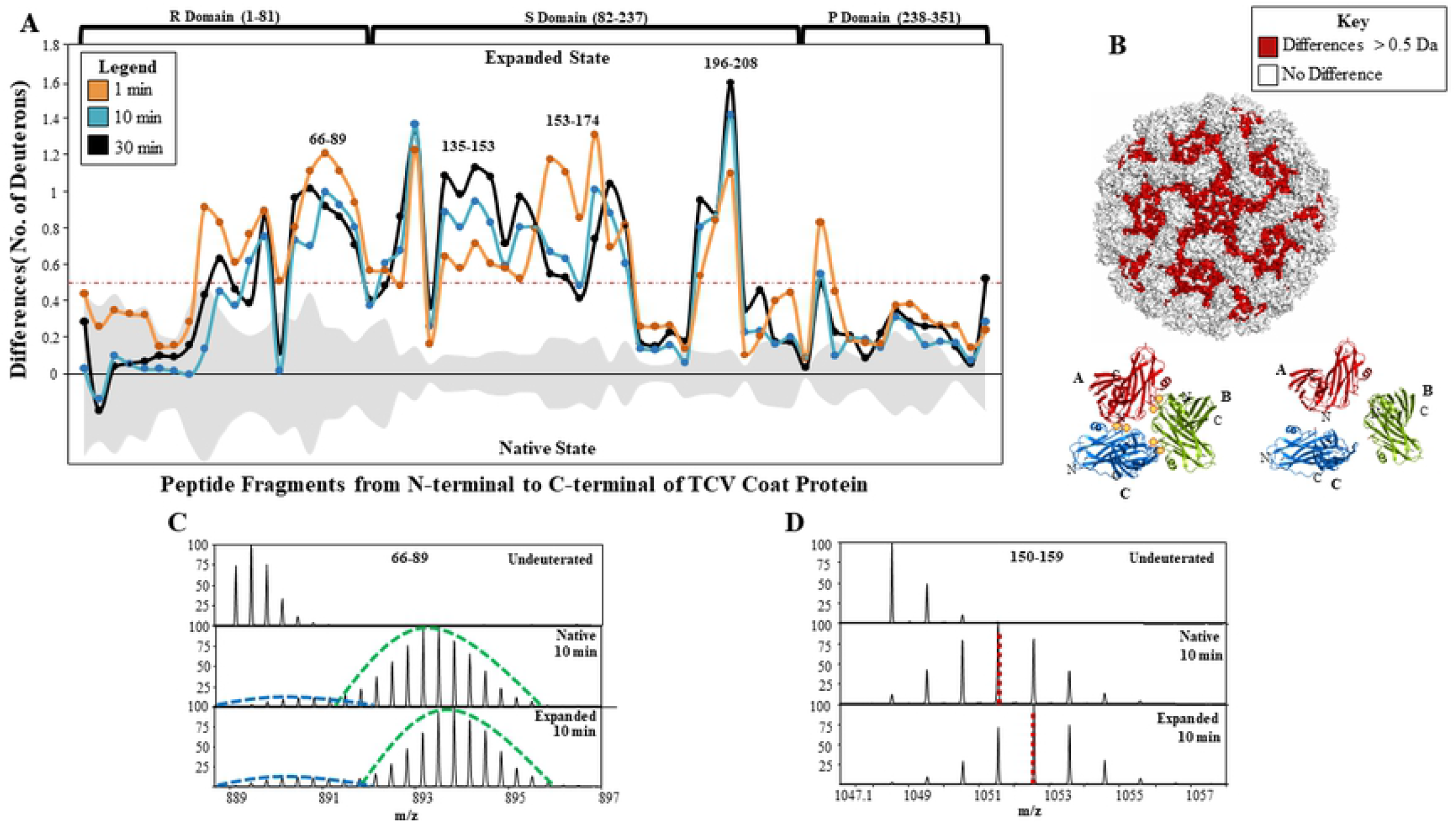
Increased proportion of low exchanging R-domain conformation in the expanded state. **(A)** Difference plot of absolute number of deuterons exchanged between expanded and native states of TCV. Each node represents pepsin proteolyzed peptides spanning TCV capsid protein after 1 (orange), 10 (blue) and 30 min (black) of deuteration respectively. **(B)** Deuterium uptake differences mapped on to TCV virus structure (PDB ID - 3ZX8) in surface representation where the red regions represent those exhibiting exchange greater than the 0.5 Da threshold showing expansion hotspots, and white representing regions showing no change following 10 minutes of deuteration. Cartoon representation of Native and Expanded state of the quasi 3-fold axis. A, B and C conformations are colored in red, green and blue respectively. Bound calcium ions at homology-based calcium binding sites (E127, D155, D157 and D199) are shown as yellow circles. **(C)** Mass spectral plot of R domain peptide 66-89 comparing native and expanded states at Dex = 10 min. The low exchanging population is outlined in blue and the high exchanging population is outlined in green. **(D)** Mass spectral plot of 3-fold axis representative peptide 150 – 159 from the S domain comparing native and expanded states at Dex = 10 mins.

The R domain peptides once again showed a characteristic bimodal exchange profile as seen in native TCV (Fig 3C). Deconvolution of mass spectra indicated a larger percentage of the lower exchanging population (~13.6%) compared to native TCV for peptide 66-89 (S2 Fig). Further, the average deuterium exchange observed for this population was much lower (~1.8 Da) in expanded relative to native TCV. The higher exchanging population in both states showed comparable average deuterium exchange (~12.7 Da) (S2 Fig).

### Expansion elicits conformational rearrangements in RNP

There are two possible explanations for the bimodal isotopic envelopes observed within the mass spectra of deuterated peptides from the R domain. Either the R domain populates a more folded and ordered conformation (low exchanging), and a disordered conformation (high exchanging) which would be consistent with EX1 kinetics of HD exchange [20], or the two populations reflect a differential effect of the RNA genome on the R domain. To identify the two populations of TCV capsid protein R domain in solution, we disassembled TCV by chelating Ca^2+^ through addition of EDTA and introducing high osmolality (>500 mM NaCl). SDS-PAGE gels of limited chymotrypsin proteolysis of TCV in the presence of 250/500 mM NaCl indicated complete cleavage of the 38 kDa coat protein to the 30 kDa cleavage product confirming complete disassembly (S3 Fig) [21,22]. We next resolved the two conformations with size-exclusion chromatography (SEC) broadly into two fractions consisting of RNA bound TCV coat protein (RNA-capsid core-complex) and the RNA-free TCV coat protein (Fig 4A) as described [21]. This allowed comparative HDXMS of a disassembled and chromatographically resolved higher molecular weight fraction containing genomic RNA-capsid core-complex alone, disassembled TCV, and native virus (Fig 4B).

**Figure 4.**
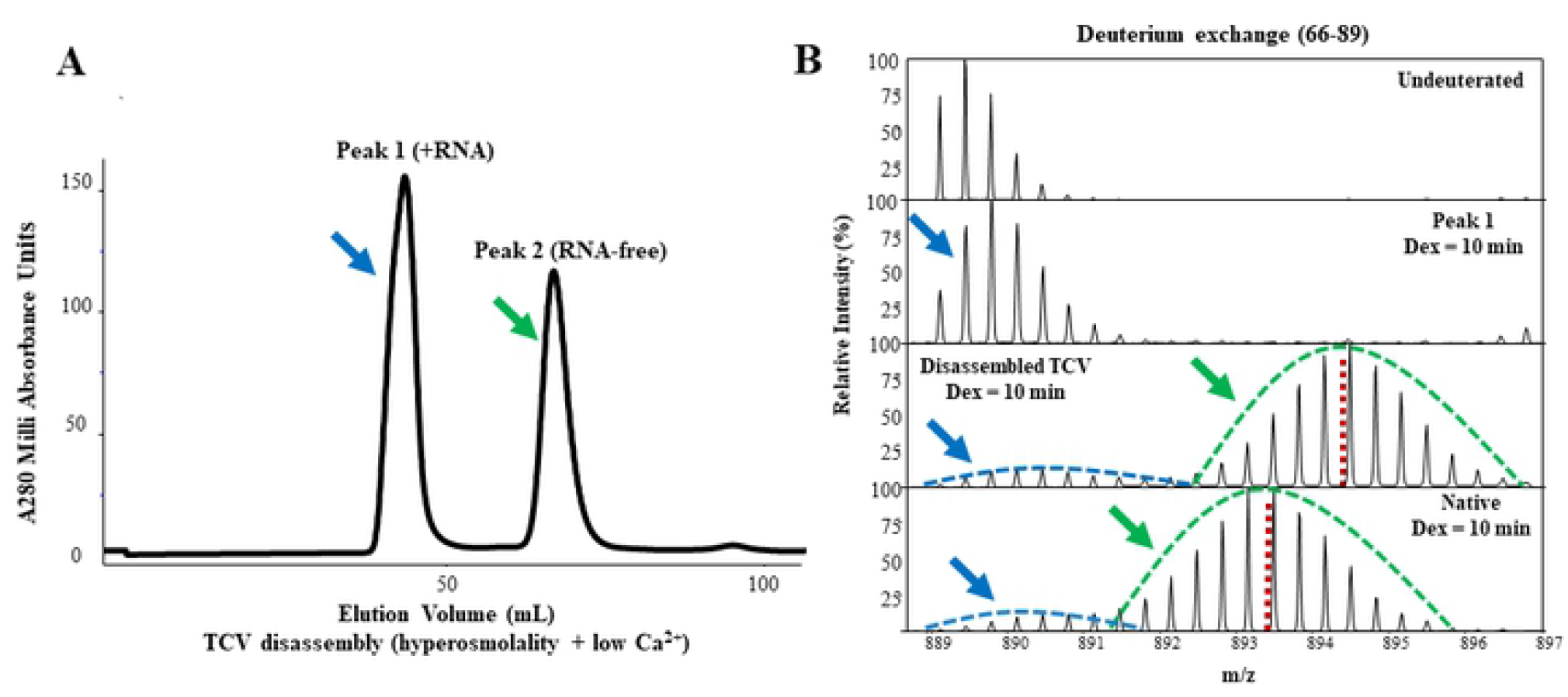
The native virus particle is comprised of low exchanging RNA-bound and free conformations. **(A)** Size exclusion chromatography of TCV disassembled in the absence of Ca^2+^ and 500 mM NaCl, indicating two well resolved peaks [21] - a fraction containing genomic RNA (RNA-capsid core complex) eluted first (Peak 1) followed by an RNA-free R-protein at a later elution volume (Peak 2). **(B)** Comparison of spectral plots of peptide 66-89 between three states-The RNA-capsid core-complex, Native state and disassembled state at Dex = 10 min. The blue arrow in the spectral plot of the RNA-capsid core-complex corresponds to the low deuterium exchanging population. The green arrow represents the free R domain conformation.

The RNA-bound fraction showed low deuterium exchange for the R domain peptides while the RNA-free fraction corresponded to the higher exchanging population. Figure 4B shows the mass spectra of deuterium exchanged peptide (66-89) for the two fractions in the disassembled virus in comparison to native TCV. The isotopic envelope of the RNA bound population (Peak 1) can be seen corresponding to the lower exchanging envelope in the corresponding native virus. TCV incubated with 500 mM NaCl contains a mixture of both high and low exchanging populations (Peaks 1 and 2) and the low exchanging population corresponds to the lower exchange in the RNA-capsid core-complex implying that the low exchanging population in the native virus particle represented the proportion of the R domain tightly bound to genomic RNA.

Importantly, the higher exchanging population of the native state showed lower deuterium exchange compared to free coat protein from the disassembled virus (Fig 4B). This indicated that the higher exchanging population represented the fraction of R domain peripherally bound to the genomic RNA in the native particle, which is disrupted by high salt, leading to disassembly of TCV into the RNA-capsid core-complex and free TCV coat proteins. While mass spectra of all the deuterium exchanged R domain peptides allowed an estimate (~13.6%) of the relative abundance of RNA-bound and free coat protein, the mass spectra of deuterium exchanged peptide 66-89 alone allowed the most accurate quantitation to a baseline resolution estimate of the relative abundance of RNA-bound and RNA-free TCV coat protein (S2 Table).

### Structure of Expanded TCV at 3.2 Å resolution shows RNA primed for release at 5’ fold axis

In order to establish if R-domain interactions with genomic RNA were uniformly distributed across the expanded TCV particle, we solved the structure of expanded TCV, generated at pH 5.4 using molecular dynamics flexible fitting (MDFF) [23]. While a structure of expanded TCV has been previously reported [13], it has been solved at low resolution (~17Å) and offers limited insights of the interior of the virus particle containing the more dynamic core of the virus containing genomic RNA and associated capsid.

During MD simulations of the complete viral particle, an additional biasing potential based on the cryo-EM map was applied. Forces proportional to the gradient of the density map trigger the gradual transition of the structure towards the experimental map. The native TCV coordinates solved at pH 7.4 (PDB: 3ZX8) were used as the starting point [13]. Initial fitting with the TCV at pH 5.4 density map determined at ~3.2 Å resulted in a correlation of ~0.41 (see Methods for details). We next trimmed the low pH map around the TCV trimeric subunit made of the 3 quasi equivalent conformations A, B and C in their native state, and then applied MDFF to progressively fit the structure into the map over 50 ns of simulation (Fig 5). Following the MDFF procedure, the backbone root mean square deviation (RMSD) of the trimer was measured with respect to the final expanded coordinates, without least-squares fitting, revealing a gradual reduction in RMSD from ~7 to 0 Å (S4A Fig). This reflected an overall translational motion of the trimer during radial expansion of the capsid. However, the trimer retained its native structure and interface; when performing a least-squares fit of the final conformation compared to the initial structure, a lower backbone RMSD of ~3.2 Å was measured, thus retaining the close proximity of key acidic residues at the quasi 3-fold axis that coordinate Ca^2+^ (S4B Fig).

**Figure 5.**
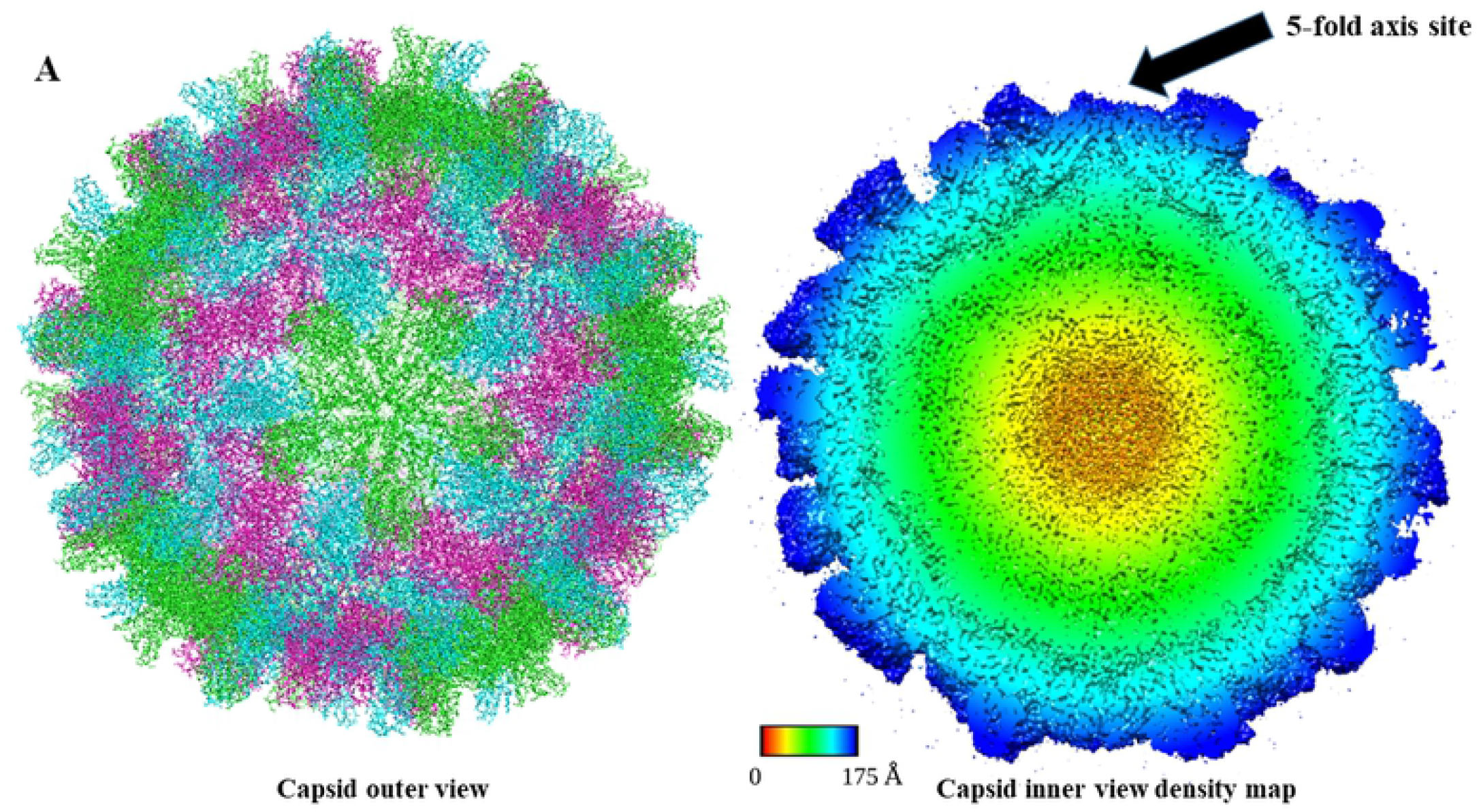
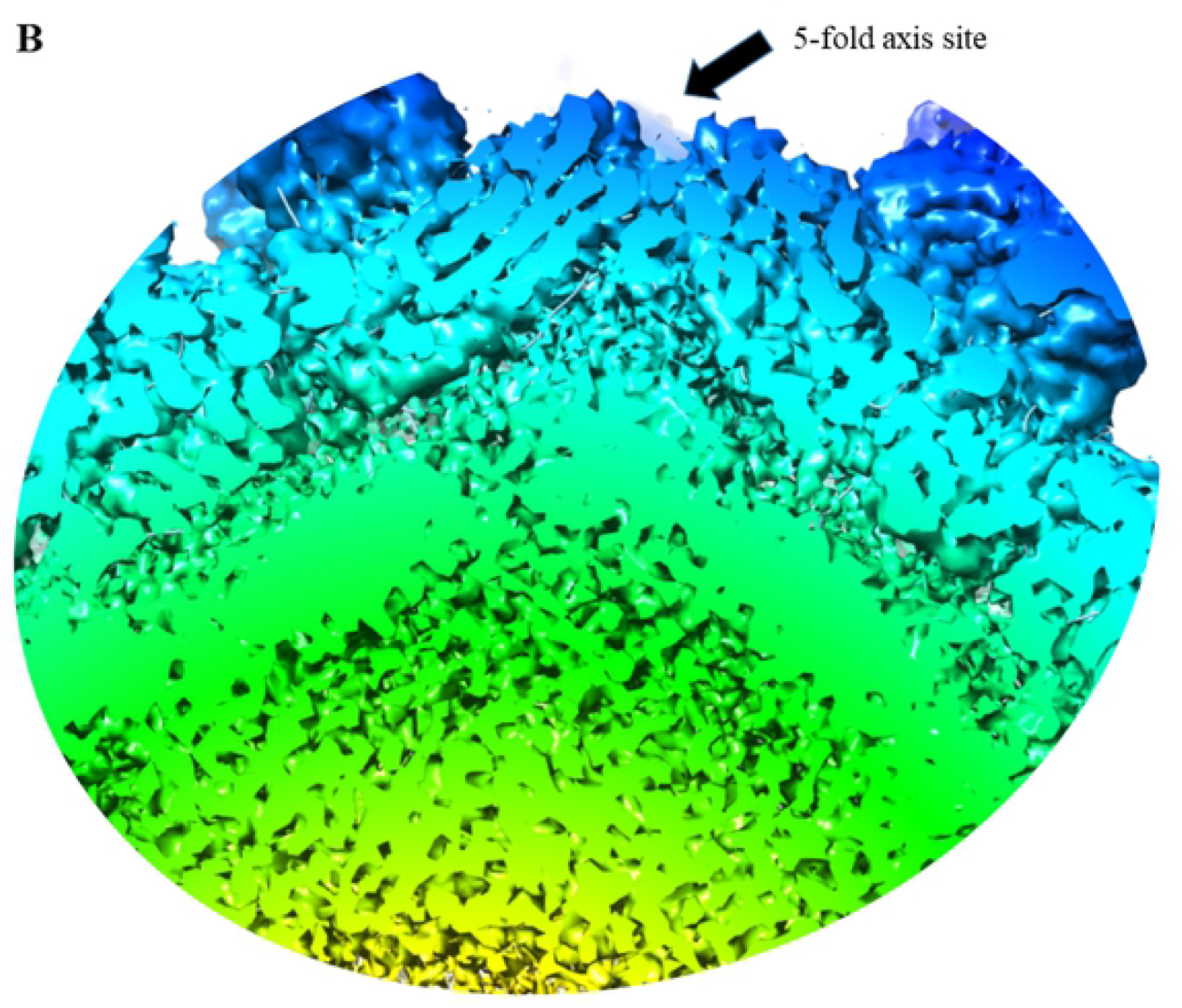
Structure of the pH 5.4 Expanded TCV capsid at 3.2 Å resolution reveals asymmetry in RNA egress. **(A)** The reconstruction of the capsid protein shell based on biased molecular dynamics flexible fitting (MDFF) of a single trimeric asymmetric unit at pH 7.4. The final snapshot of the trimer was subsequently aligned on the low pH coordinates (PDB: 3ZX8). The TCV capsid is colored according to the asymmetric unit (A-red, B-green, C-blue) shown in Figure 1, panel B. **(B)** The cross section of the Cryo-EM density map is colored according to its radius as shown inset. The final correlation of the expanded TCV structure and the cryo-EM density map resulted in ~0.80.

The final fitted trimeric coordinates were aligned onto the 60 subunits of the initial TCV model. This resulted in an accurate model for the expanded state of the virus, as evidenced by a correlation measured with respect to the low pH density map of ~0.80, representing a significant improvement in comparison to the initial fit (~0.41). The final coordinates together with the corresponding cryo-EM density map of the expanded TCV at low pH are shown (Fig 5). This reveals not only a larger diameter of the capsid when compared to the native structure, but interestingly asymmetric packing at the 5-fold axes with the pentamers protruding outwards from the TCV. Importantly, the protrusion point is likely the single copy of the covalently bound p80 dimer. The p80 dimer is preferentially labelled by I^125^, showing it to be on the surface. It is additionally part of the RNA-bound core protein, implicated as the last segment of RNA to undergo condensation during assembly and correspondingly, the first RNA-coat protein segment to be released from the viral particle [24]. This reveals the asymmetric packing of RNA within the TCV virion and allows visualization of a viral RNA genome in the process of release. Even at a 3.2 Å resolution we were unable to capture the structure of the RNA within the particle indicating that the genomic RNA in the interior of the particle is highly dynamic. The increased proportion of RNA-bound R domain can be attributed to enhanced interactions mediated by protruding RNA at the 5’ fold site with proximal R domain.

### A stable RNP complex is retained upon complete TCV disassembly in the presence of osmolyte

The expanded particle undergoes full disassembly upon addition of osmolyte (Fig 4). To map changes accompanying disassembly, we carried out HDXMS in three osmolyte (NaCl) concentrations (100 mM, 250 mM, and 500 mM). We observed increased exchange in the high exchanging population of the R domain (peptide 66-89 in S5A Fig) and at the 5-fold/3fold axis (peptide 135-154 in S5B Fig) corresponding to particle disassembly. At 250 mM NaCl the TCV particle showed nearly complete disassembly as shown in the difference plot in S5C Figure. Interestingly, the low exchanging RNA bound fraction of R domain did not show significantly increased exchange under high osmolyte conditions, suggesting that the RNP complex is maintained upon complete disassembly (S5A Fig).

## DISCUSSION

Viruses are evolutionarily built to encounter multiple environments within and outside hosts during their lifecycle. Abiotic changes such as temperature, pH, osmolyte, and divalent cations can all serve as indicators for the virus while seeking favorable conditions for disassembly. RNA viruses including Dengue, flaviviruses and the common cold undergo dynamic breathing motions in order to modulate behavior under these changing conditions and to help evade host defences [19]. A holistic view is needed to account for the contributions of the structure of the viral particle as well the dynamic changes that occur in the disordered interior. We have uniquely carried out doing dynamics measurements with HDXMS in conjunction with orthogonal cryo-EM. This combination allows for a complete view of viral response to environmental perturbations.

The RNA-protein interface is a critical and poorly understood component of viral dynamics. TCV has no lipids and no glycosylation modifications offering a deeper view into the role of RNA-protein interactions in viral disassembly. The simplicity of TCV means that the viral capsid protein must have an inbuilt efficiency in order to function in many roles including as a scaffold for genomic RNA and as a sensor for environmental perturbations. We have outlined the changes in TCV capsid protein during disassembly where the particle responds to changes in divalent cations and osmolyte in host plant cells during disassembly.

Viruses are highly dynamic biomolecular entities undergoing molecular motions of varying timescales that facilitate replication within host cells. This study has offered key insights into the virus interior that is structurally poorly resolved and has revealed the key sensors of viral metastability at the RNA-R domain interface. From cryo-EM, it was predicted that genomic RNA mediates strong interactions with ~30% of the coat protein R domains, potentially correlated with the formation of the folded arm of the C quasi equivalent conformation [21]. Our results reveal that the R domain provides the main interface for RNA binding and presents a more accurate smaller percentage of 5.7% being RNA-bound in native TCV. This represents the RNA-bound capsid core of TCV. The remaining relatively free coat protein in the (~94.3%) fraction is loosely bound to the RNA-capsid protein core in native TCV (Fig 6).

**Figure 6.**
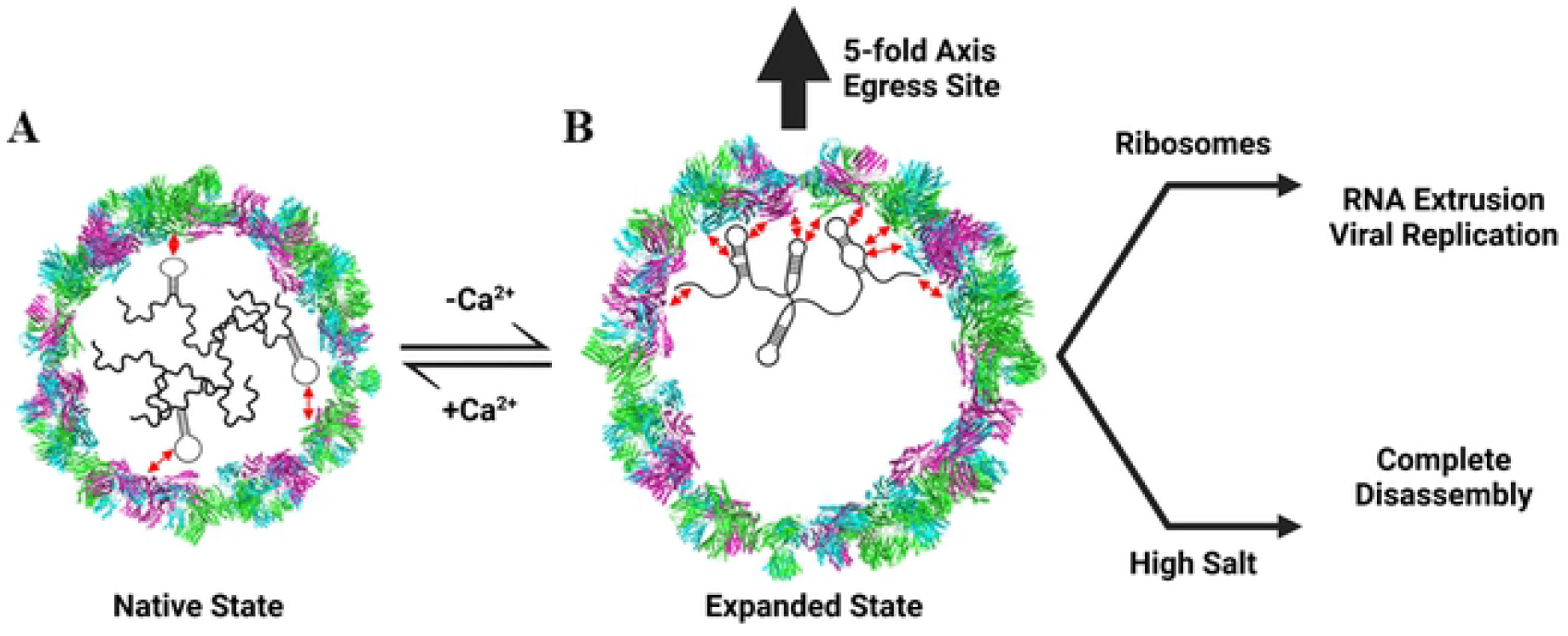
TCV disassembly-assembly model: Viral breathing and RNP complex drives programmed TCV disassembly. **(A)** Cross-section of TCV in the native state (PDB: 3ZX8, Diameter = 300 Å) shown encapsidating genomic RNA (black tangled line). RNA interactions with the capsid protein (blue, pink, and green monomers) are shown as red double-sided arrows. **(B)** Upon calcium depletion (either in the plant host cell or by addition of EDTA), the TCV particle expands. The genomic RNA is represented as more ordered with additional contacts between RNA and capsid shown. The black arrow demonstrates RNA egress from an asymmetric five-fold axis point. The RNA can be extruded by ribosomes in host cells or particle disassembly can be induced under high osmolyte conditions.

These results reveal the critical importance of the genomic RNA capsid protein complex to function as a sensor for the host environment. Increased osmolyte and decreased Ca^2+^ in the host environment disrupts weak contacts between the capsid and RNA genome during expansion. Conformational changes in the genomic RNA-capsid core lead to the detection of a major (~86.4%) unbound population of R domain, and a minor (~15%) strongly bound population of R domain in the expanded particle. This conformational change primes the RNA for future release. Release can then occur *in vitro* in the presence of high osmolality (500 mM NaCl) which leads to continued association of the RNA-capsid core. *In vivo*, the primed RNA genome is extruded from the expanded particle by cellular ribosomes as was shown previously [13].

Our cryo-EM analysis has revealed that the genomic RNA-capsid interactions are not equally distributed throughout the expanded particle. Instead, an asymmetric packaging was observed with a protrusion point a one specific 5-fold axis that also likely represents the covalent p80 dimer. Additionally, increased density from the interior of the particle can be seen near this protrusion point. The covalent dimer is also part of the genome capsid core during assembly, suggesting that the genomic RNA acts as a nucleation point for the RNA capsid core during assembly. This RNA-capsid core is then maintained in the whole viral particle and mediates viral metastability and environmental sensing. During the disassembly process, the RNA directs extrusion toward one specific 5-fold axis.

These results underscore the significance of RNA structure and conformational dynamics of RNA-capsid cores in viral sensing and disassembly. We have tracked dynamics of TCV through the entire particle disassembly process (Fig 6). During disassembly, the TCV particle goes through an expanded intermediate state upon Ca^2+^ depletions. TCV particle expansion also corresponds to an increase in the fraction of tightly bound RNA-R domain and conformational changes in the RNA-capsid core. Under high osmolyte conditions, the virion is disassembled into the RNA-capsid core (15% of the total coat protein) and free coat protein. In the host cell environment, ribosomes along with protease activity extrude the emerging RNA during disassembly [13]. The principles governing how RNA self assembles and disassembles upon sensing the optimal environmental cues are still to be understood. However, it is clear that the genomic RNA is in fact not a passively packaged entity but instead plays a key role throughout the TCV viral lifecycle as an environmental sensor and a driving force behind viral disassembly.

## MATERIALS AND METHODS

### Purification of virus particles and size exclusion chromatography

Native TCV particle purification was adapted based on a protocol described previously [25]. About 100 g of infected leaves were homogenized in a cold blender with 300 mL pre-chilled extraction buffer (0.2 M NaOAc, 50 mM NaCl, 20 mM CaCl_2_, 5 mM EDTA, 0.1% 2-Mercaptoethanol pH 5.4) for 10 min, with a 30 s pause for every minute. The resulting slurry was centrifuged for 15 min at 9000 rpm at 4 °C using a JA14 rotor (Beckman Coulter, Brea, CA). The supernatant was filtered with a Miracloth and kept on ice. Equal volumes of supernatant and ammonium sulfate were mixed and incubated at 4° C for 3 hours followed by centrifugation at 9000 rpm at 4 °C (JA14). The pellets obtained were resuspended in a total of 50 mL of resuspension buffer (50 mM NaOAc, 50 mM NaCl, 20 mM CaCl_2_, 0.1% 2-Mercaptoethanol, 1% Triton-X pH 5.4) and left to gently shake overnight at 4 °C. The virus was pelleted by centrifugation at 9000 rpm with JA14 at 4 °C and the supernatant was transferred to an SW28 tube. 4 mL of 10% (w/v) sucrose was transferred to the bottom of the tube using a Pasteur pipette and centrifuged at 27,000 rpm for 3 hours at 4° C using an SW28 Ti rotor (Beckman Coulter, Brea, CA). Each pellet containing the virus particles was resuspended in 1 mL resuspension buffer overnight with gentle shaking at 4 °C. All purification steps were performed at 4°C.

Resuspended particles were layered onto a 15-45% (w/v) cesium chloride continuous density gradient in resuspension buffer and centrifuged at 34,000 m for 5 hours at 20 °C using an SW41 Ti rotor (Beckman Coulter, Brea, CA). The virus particles were observed in the lower of two pale white bands in the gradient; these were extracted and dialyzed against 25 mM Tris pH 7.5 at 4 °C and measured using a Nanodrop One (Thermo Fisher Scientific, Waltham, MA). Particles for cryo-EM were dialyzed against 20 mM NaOAc, pH 5.4 at 4 °C. ~5 mg TCV native particles were incubated overnight at a final concentration of 25 mM Tris, 5 mM EDTA, 500 mM NaCl pH 7.5 (dissociation buffer). The resulting dissociated particles were subjected to size exclusion chromatography using a Superdex^™^ 200 16/60 GL column pre-equilibrated with 2 column volumes of dissociation buffer, in an AKTA^™^ FPLC system (General Electric Healthcare, Chicago, IL). Peaks 1 and 2 were collected and identified as the RNA-capsid core-complex and free protein as previously suggested [21].

### Genomic RNA transcription

cDNA stretches corresponding to the 3’ UTR region of TCV genomic RNA (5’-CAACUGAGGAGCAGCCAAAGGGUAAAUUGCAAGCACUCAGAAU-3’) were obtained from GenScript [26]. ssRNA transcription was performed using Ambion MEGAscript RNAi Kit obtaining ~2500 ng/μl of pure RNA. TCV coat protein was incubated with TCV RNA for 30 min on ice at a molar ratio of 1:1 prior to the deuterium exchange reaction [22]

### Amide Hydrogen Deuterium Exchange

Virus samples were treated with buffers as mentioned: Native (25 mM Tris, pH 7.5), NaCl titration (25 mM Tris, 5 mM EDTA, 0.1/0.25/0.5 mM NaCl, pH 7.5) to monitor conformations of native, Ca^2+^-depleted and in increasing osmolyte environments. All samples were diluted to a final concentration of 93.3% D_2_O to initiate the deuterium exchange reaction. Deuterium buffers were prepared by desiccation of respective aqueous buffers and reconstituted in equivalent volumes of D_2_O. Deuterium exchange was carried out at room temperature (26 °C) maintained on a drybath for 1, 10 and 30 min followed by rapidly quenching the reaction to minimize back exchange using 4 M GdnHCl and 0.1% TFA on ice to bring the pH down to 2.5.

### Pepsin proteolysis and mass spectrometry analysis

Quenched samples were injected onto an immobilized pepsin treatment (BEH Pepsin Column, Enzymate, Waters, Milford, MA) using a nano-UPLC sample manager at a constant flow rate of 100 μl/min of 0.1% formic acid. Proteolyzed peptides were then trapped in a VanGuard column (ACQUITY BEH C18 VanGuard Pre-column, 1.7 μm, Waters, Milford, MA) and separated using a reversed phase liquid chromatography column (ACQUITY UPLC BEH C18 Column, 1.0 × 100 mm, 1.7 μm, Waters, Milford MA). NanoACQUITY binary solvent manager (Waters, Milford, MA) was used to pump an 8-40% acetonitrile gradient at pH 2.5 with 0.1% formic acid at a flow rate of 40 μl/min and analyzed on a SYNAPT G2-S_i_ mass spectrometer (Waters, Milford, MA) acquired in MS^E^ mode [6].

Undeuterated TCV particles were sequenced by MS^E^ to identify pepsin digested peptides using Protein Lynx Global Server Software (PLGS v3.0) (Waters, Milford, MA). The peptides were identified by searching against the TCV coat protein sequence database (UniProt ID: P06663) with a non-specific proteolysis enzyme selected. Peptides from the undeuterated samples that were identified and matched from the primary sequence database were filtered and considered with the following specifications: precursor ion tolerance of < 10 ppm, products per amino acid of at least 0.2 and a minimum intensity of 1000.

Average deuterium exchange in each peptide was measured relative to undeuterated control peptides using DynamX v3.0 (Waters, Milford, MA) by determining the centroid mass of each isotopic envelope. Subtractions of these centroids for each peptide from the undeuterated centroid determined the average number of deuterons exchanged in each peptide [18]. For bimodal populations, centroid masses of each individual population were assessed independently. Percentage of each exchanging conformation was calculated as the ratio of intensity of ion sticks corresponding to each conformation divided to the total intensity sum of all ion sticks associated with deuterium exchange for that peptide.

Deuterium exchange for all peptides is represented using relative fractional uptake (RFU) plots. Each value reported is an average of three independent deuterium exchange experiments and not corrected for back-exchange [6]. Difference plots were made by subtracting absolute centroid mass values between the two states under consideration. A difference of ± 0.5 Da was considered a significance threshold for deuterium exchange [27].

### Deconvolution analysis of deuterium exchange mass spectral envelopes

Deconvolution of mass spectral envelopes was carried out by HDExaminer version 3.2 (Sierra Analytics, Modesto CA) to check for ensemble behavior in solution, specifically to check for two distinct distributions of mass spectral envelopes [19]. Each of these envelopes would represent distinct conformations in solution [20]. We set a threshold fit value score of 0.9 for the goodness of fit of the experimental mass spectral envelope for deuterium exchanged peptides with the theoretical envelope. If the fit score for a peptide is less than the threshold, the program will fit the spectral envelope to a bimodal distribution. If the score of the bimodal distribution is greater than that of the unimodal distribution, the program assigns a lower exchanging (left population) and a higher exchanging (right population). Results from deconvolution for mass spectra (peptide 66-89) for each of the three states are shown in S. Figure 2 and S. Table 2 and 3.

### Limited chymotrypsin proteolysis

Native, expanded and disassembled TCV particles (native and increasing concentrations of NaCl) were treated with Chymotrypsin for 10 min at a ratio of 1:100 of protease-protein (w/w) at room temperature as described previously [13].

### Cryo-electron microscopy

5 μl (2 mg/ml) of TCV purified virions suspended in 20 mM NaOAc (pH 5.4) were applied onto the holey carbon grid (Quantfoil 2/2 200 mesh) after 1 minute glow-discharge, subsequently the grid was blotted and plunge frozen with Vitrobot Mark IV(Thermo Fisher Scientific, USA). The grids were loaded into Titan Krios Cryo-electron microscope (Thermo Fisher Scientific, USA) and more than 3000 micrographs was collected at nominal magnification of 96 kX (pixel size 0.89 Å) with Falcon-II CMOS electron detector (Thermo Fisher Scientific). A total of 19353 virus particles were picked manually, and the particle set was processed with Frealign v9.09 for 3D reconstruction and refinement. The final resolution reached 3.2 Å.

### Molecular dynamics flexible fitting (MDFF)

The UCSF Chimera software was used to trim the low pH map around the TCV trimer as well as to fit and calculate the correlation of the atomic structure into the cryo-EM density map [28]. The correlation between fitted coordinates and the density map varies from −1 to +1, from decorrelated to identical, respectively. The *Fit in Map* function in Chimera locally optimizes the fit of atomic coordinates into the density map using translation and rotation by maximizing their overlap.

Each of the monomers of TCV corresponded to residue numbers 53-351, where the first 52 residues corresponded to the unresolved, unstructured N-terminal region. The trimeric construct in its native state was placed in a rectangular box of dimensions ~13×15×13 nm^3^ and solvated with ~80,000 explicit TIP3P water molecules [29]. Protein parameters corresponded to the CHARMM36m force field [30]. Chloride ions were used to neutralize the overall system charge. The total system size corresponded to ~260,000 atoms. The energy of the system was minimized with 25,000 steps using the conjugate gradient algorithm. Equilibration in the *NVT* ensemble was performed for 1 ns while applying position restraints on all protein heavy atoms with a force constant of ~20 kcal mol^-1^ Å^-2^. Equations of motion were integrated through the velocity Verlet algorithm with a 1 fs time step together with Langevin dynamics to maintain the temperature at 310 K [31]. A cutoff distance of 1.4 nm was used for the short-range neighbor list and for van der Waals interactions. The Particle Mesh Ewald method was applied for long-range electrostatic interactions with a 1.2 nm real space cutoff [32]. MDFF was run in 25 steps of 2 ns each, totalling 50 ns of sampling. Each step of MDFF employed a progressive increase in the factor of the grid scale potential (from 0.3 to 15). All simulations were performed using NAMD software [33] on: i) an in-house Linux cluster composed of 8 nodes containing 2 GPUs (Nvidia GeForce RTX 2080 Ti) and 24 CPUs (Intel^®^ Xeon^®^ Gold 5118 CPU @ 2.3 GHz) each; and ii) ASPIRE 1, the petascale cluster at the Singapore National Supercomputing Centre, where each simulation employed 4 nodes each consisting of 1 GPU (Nvidia Tesla K40t) and 24 CPUs (Intel^®^ Xeon^®^ CPU E5-2690 v3 @ 2.6 GHz). Visualization of simulation snapshots used the Chimera software.

## ACKNOWLEDGEMENTS

This work was initiated by a grant from Singapore Ministry of Education Academic research fund – Tier 1 [MOE2018-T1-A73-114] awarded to G.S.A and further supported by startup funds from The Pennsylvania State University to G.S.A.

## SUPPLEMENTARY FIGURE LEGENDS

**Supplementary Figure 1. Primary sequence coverage map of pepsin proteolyzed peptides of TCV coat protein unit.** Coverage map showing 61 Peptides spanning the TCV coat protein unit - 19 in the R-domain (orange) spanning 1 – 81 out of which 5 peptides overlap the S domains (blue), 30 in the S-domain spanning 82 - 238 and 12 in the P-domain (green) spanning 239 - 351 with total coverage being 89.2% and 2.87% redundancy. The limited proteolysis site Tyrosine 66 (Y66) is marked in red. All peptides spanning this region are marked with a red bar in region corresponding to Y66.

**Supplementary Figure 2. HDEXaminer deconvolution of states in peptide 66-89. (A)** Deconvolved spectra of TCV peptide 66-89 in the native state. The low exchanging population (yellow) accounts for 5.7% of native state R-domain peptides with an uptake of 6.2 Da. The high exchanging population (red) accounts for 94.3% of native state R-domain peptides with an uptake of 11.7 Da. **(B)** Deconvolved spectra of TCV peptide 66-89 in the expanded state. The low exchanging population (yellow) accounts for 13.6% of expanded state R domain peptides with an uptake of 1.8 Da. The high exchanging population (red) accounts for 86.4% of R domain peptides with an uptake of 12.7 Da.

**Supplementary Figure 3. The rate of limited proteolysis is dependent on osmolyte concentration.** SDS-PAGE of limited chymotrypsin proteolysis on TCV coat protein monomer (10 min digestion, 1:100 w/w of protease and TCV coat protein) for native and NaCl induced states – 100 mM, 250 mM and 500 mM. The 38 kDa band corresponds to the whole TCV coat protein unit while the 30 kDa bands have been shown to be a cleavage product consisting of the S and P domains through N-terminal sequencing [34].

**Supplementary Figure 4. RMSD of the expanded TCV trimeric axis. (A)** Root mean square deviation (RMSD) plot of the 3-fold axis of the TCV trimer (black) with respect to coordinates of expanded TCV (black). X-axis represents RMSD in Å and Y-axis represents the simulation time in ns. **(B)** Cartoon representation overlaid on the surface density of the asymmetric trimeric unit of TCV in the native state bound to calcium ions (yellow spheres). The A, B and C monomers of TCV are colored in red, green and blue, respectively. In the left is a cross-sectional view and is rotated 90° and represented as a longitudinal of the asymmetric trimer.

**Supplementary Figure 5. Addition of NaCl leads to particle disassembly** Comparison of mass spectral plots of **(A)** R-domain peptide 66-89 and **(B)** S-domain peptide 135-154 from TCV in the presence of 100 mM, 250 mM and 500 mM NaCl. The low exchanging population for peptide 66-89 is outlined in blue and the high exchanging population is outlined in green. **(C)** Deuterium exchange difference plot of TCV with 100 mM (blue), 250 mM (green) and 500 mM NaCl (orange) relative to the native state at Dex = 10 min. Regions shaded light pink indicate peptides showing large increases in exchange with NaCl. Listed on X-axis are pepsin-fragment peptides listed from N to C-terminus. Y-axis corresponds to the absolute differences in deuterium exchanged. Dotted red lines are indicative of the 0.5 D deuterium exchange difference threshold while the standard error is represented in grey.

## SUPPLEMENTARY TABLES

**Supplementary Table 1. Table of fragment peptides and ion products obtained from TCV Native virus after acquiring by MS^E^ mode.** Ion product fragmentation profiles of all the 61 peptides of the TCV coat protein in the native state.

